# Association between Cortical Thickness and Functional Response to Linguistic Processing in the Occipital Cortex of Early Blind Individuals

**DOI:** 10.1101/2025.04.17.645592

**Authors:** Maria Czarnecka, Katarzyna Hryniewiecka, Agata Wolna, Clemens Baumbach, Joanna Beck, Jyothirmayi Vadlamudi, Olivier Collignon, Katarzyna Jednoróg, Marcin Szwed, Anna-Lena Stroh

## Abstract

Blindness has been shown to induce changes in the structural and functional organization of the brain. However, few studies have investigated the relationship between these structural and functional changes. In this study, we examined cortical thickness within occipital regions of interest in 38 early blind individuals and explored its relationship to functional activation during linguistic processing. Participants engaged in tactile Braille reading and auditory processing tasks involving words, pseudowords, and control conditions to assess various aspects of linguistic processing. Linear mixed models revealed a significant association between cortical thickness and functional activation in the occipital cortex during linguistic tasks. Specifically, lower cortical thickness in the middle occipital gyrus, the calcarine sulcus, and the parieto-occipital sulcus were linked to increased activation during orthographic processing in blind participants (Braille pseudowords vs. Braille nonsense-symbols). Similarly, lower cortical thickness in the calcarine sulcus and parieto-occipital sulcus was associated with greater functional activation during phonological processing (auditory pseudowords vs. auditory control). These findings align with prior research suggesting that structural and functional adaptations in the visual cortex of blind individuals may be influenced by developmental mechanisms such as pruning or myelination. This study highlights the interplay between cortical structure and functional reorganization in the blind brain.

## INTRODUCTION

It is well-established that the functional and structural organization of occipital cortices of blind individuals differs drastically from that of sighted individuals. For example, regions that are typically associated with visual processing in sighted individuals, have been shown to be activated during both tactile and auditory processing in blind individuals (Bedny et al., 2011; Collignon et al., 2011; Dzięgiel-Fivet et al., 2021; Röder et al., 2002; Sadato et al., 1996). Early blindness also triggers changes in structure, like an increase in cortical thickness within the occipital cortex (Aguirre et al., 2016; Anurova et al., 2015; Bridge et al., 2009; Hasson et al., 2016; Jiang et al., 2009; Park et al., 2009; Voss & Zatorre, 2012). However, it is unclear whether these structural and functional changes are linked.

Numerous studies have reported that blind individuals show enhanced recruitment of occipital cortices when processing non-visual information. Interestingly, this crossmodal recruitment of the occipital cortex of blind people seems to follow similar organizational principles as the occipital cortex of sighted individuals (Beisteiner et al., 2015; Dzięgiel-Fivet et al., 2021; Rączy et al., 2019; Reich et al., 2011; Sadato et al., 1996). For instance, parts of the ventral occipito-temporal cortex (vOTC) that have been associated with reading in sighted individuals, during Braille reading, or the recruitment of the visual “where” pathway during sound localization(Collignon et al., 2011; Gougoux et al., 2004), as well as the activation of the visual fusiform face area during voice identity processing (Hölig et al., 2014; Mattioni et al., 2020).

In addition to these functional changes, blindness has been shown to trigger several structural brain changes. Arguably the most documented observation is that blind individuals seem to have thicker occipital cortices compared to sighted individuals (Bridge et al., 2009; Hölig et al., 2014; Jiang et al., 2009; Park et al., 2009). The mechanisms causing the apparent increase in cortical thickness are still a matter of debate. While Jiang and colleagues proposed that increased cortical thickness in blind individuals is a result of a disruption in synaptic pruning (Jiang et al., 2009), Park and colleagues instead hypothesized that it was the result of cross-modal plasticity (Park et al., 2009). However, these two proposals are not mutually exclusive.

On one hand, it could be that cross-modal recruitment of occipital areas is possible because certain cortico-cortical connections are not pruned, but rather preserved and strengthened (Singh et al., 2018). Indirect evidence for this notion was provided by Voss and Zatorre, who observed that the cortical thickness of occipital areas was related to behavioral enhancements in pitch and melody discrimination in blind individuals (Voss & Zatorre, 2012). Similarly, Stroh et al. have observed that blind individuals with thicker occipital cortices are better at sensing their own heartbeat (Stroh et al., 2024). Based on these findings, it could be hypothesized that increased cortical thickness goes hand in hand with increased functional recruitment.

However, the widely accepted idea that cortex thins during development has recently been questioned by quantitative MRI and diffusion MRI studies. These studies propose that increased myelination in the cortex alters MR image contrast influencing the perceived gray-white matter boundary (Gomez et al., 2017; Natu et al., 2019). This raises the possibility that the previously observed thicker visual cortex in blind individuals may, in fact, be due to reduced myelination in these regions (Hölig et al., 2023).

So far, only one study has assessed the relationship between structural measures and functional activation in blind individuals (Anurova et al., 2015). Authors measured cortical thickness and its relationship with cortical activity during one-back tasks for auditory localization, pitch identification, and a simple sound-detection task in early blind and sighted individuals. Functional activation during sound-localization and pitch-identification tasks correlated negatively with cortical thickness in occipital areas of early blind but not in sighted individuals (Anurova et al., 2015). They concluded that activity-dependent pruning and changes in synaptic efficacy are the primary drivers of structure-function relationships in blind individuals, though other forms of plasticity, such as those from long-term practice, may also play a role.

The aim of the current study was to further explore this structure-function relationship in a relatively large group (n=38 compared to n=12 in Anurova et al.) of blind individuals and assess whether similar relationships between structure and function can be observed for higher-level linguistic processing. Early blind participants and sighted control participants were presented with words, pseudowords, and a low-level control condition during reading (blind participants were reading tactile Braille and sighted control participants were reading visual print) and speech comprehension. Associations between cortical thickness and functional activation during reading and speech comprehension were analyzed within the occipital cortex. In contrast to previous studies, we assessed the associations between structure and function at the single-subject level by fitting the model to vertex-wise cortical thickness and percent signal change within each region of interest. Based on previous studies, we expected to observe increased activation for Braille word reading and auditory word comprehension in occipital areas, including the vOTC and early visual areas, such as the calcarine sulcus and the lingual gyrus (Bedny et al., 2011; Beisteiner et al., 2015; Sadato et al., 1996) in blind participants. Moreover, we expected to find a negative association between cortical thickness and functional activation within the occipital cortex of blind but not in sighted participants.

## METHODS

### Datasets

Data were collected at two different scanning sites using the same experimental procedure. Overall, thirty-eight early blind participants (mean age: 34.87, age SD: 9.45, 22 females) and 25 sighted participants (mean age: 35.42, age SD: 9.68, 16 females) participated in the study (see Table 2 for demographic characteristics of the blind participants). All of the blind participants declared having at most minimal light perception since birth, which did not allow for visual navigation or object or color perception. The blind participants began to learn Braille between the age of six and nine and use it daily (reading or writing). Participants declared no history of neurological illness or brain damage (other than the cause of blindness) and all of the participants declared having normal hearing. Almost all participants were right-handed (1 left-handed). Regardless of the handedness, 18 participants used the non-dominant hand for Braille reading.

**Table 1.** Demographic characteristics of the blind participants.

| Subject | Age | Sex | Cause of blindness | Handedness | Reading hand | Dataset |
| --- | --- | --- | --- | --- | --- | --- |
| 1 | 35 | F | Unknown | Right | Right | 1 |
| 2 | 36 | F | Retinopathy of prematurity | Right | Left | 1 |
| 3 | 30 | F | Retinopathy of prematurity | Right | Left | 1 |
| 4 | 44 | F | Congenital rubella syndrome | Right | Right | 1 |
| 5 | 48 | F | Unknown | Right | Left | 1 |
| 6 | 37 | F | Glaucoma | Right | Right | 1 |
| 7 | 31 | F | Leber's congenital amaurosis | Right | Right | 1 |
| 8 | 19 | F | Retinoblastoma | Right | Right | 1 |
| 9 | 28 | F | Leber's congenital amaurosis | Right | Left | 1 |
| 10 | 39 | F | Mechanical damage | Right | Right | 1 |
| 11 | 42 | F | Retinopathy of prematurity | Right | Right | 1 |
| 12 | 60 | F | Retinopathy of prematurity | Right | Right | 1 |
| 13 | 41 | F | Leber's congenital amaurosis | Right | Right | 1 |
| 14 | 24 | F | Congenital rubella syndrome | Right | Right | 1 |
| 15 | 28 | F | Optic nerve hypoplasia | Left | Right | 1 |
| 16 | 45 | F | Retinopathy of prematurity | Left | Right | 1 |
| 17 | 38 | M | Optic nerve hypoplasia | Right | Right | 1 |
| 18 | 46 | M | Optic nerve hypoplasia | Right | Left | 1 |
| 19 | 36 | M | Unknown | Right | Left | 1 |
| 20 | 21 | M | Retinopathy of prematurity | Right | Left | 1 |
| 21 | 38 | M | Retinitis pigmentosa | Right | Right | 1 |
| 22 | 52 | M | Cataract, glaucoma | Right | Left | 1 |
| 23 | 19 | M | Bilateral microphthalmia | Right | Left | 1 |
| 24 | 27 | M | Retinopathy of prematurity | Right | Right | 1 |
| 25 | 36 | M | Retinopathy of prematurity | Right | Left | 1 |
| 26 | 29 | F | Neuropathy of the optic nerve | Right | Right | 2 |
| 27 | 23 | M | Eyeball hypoplasia | Right | Left | 2 |
| 28 | 31 | M | Retinopathy of prematurity | Right | Left | 2 |
| 29 | 24 | M | Retinopathy of prematurity | Right | Left | 2 |
| 30 | 26 | M | Retinopathy of prematurity | Right | Right | 2 |
| 31 | 30 | M | Optic nerve hypoplasia | Right | Right | 2 |
| 32 | 37 | F | Retinopathy of prematurity | Right | Right | 2 |
| 33 | 44 | F | Retinopathy of prematurity | Right | Left | 2 |
| 34 | 44 | M | Retinopathy of prematurity | Right | Left | 2 |
| 35 | 39 | F | Retinopathy of prematurity | Right | Left | 2 |
| 36 | 45 | F | Retinopathy of prematurity | Right | Right | 2 |
| 37 | 27 | F | Retinopathy of prematurity | Right | Right | 2 |
| 38 | 26 | F | Retinopathy of prematurity | Right | Left | 2 |

**Table 2.**
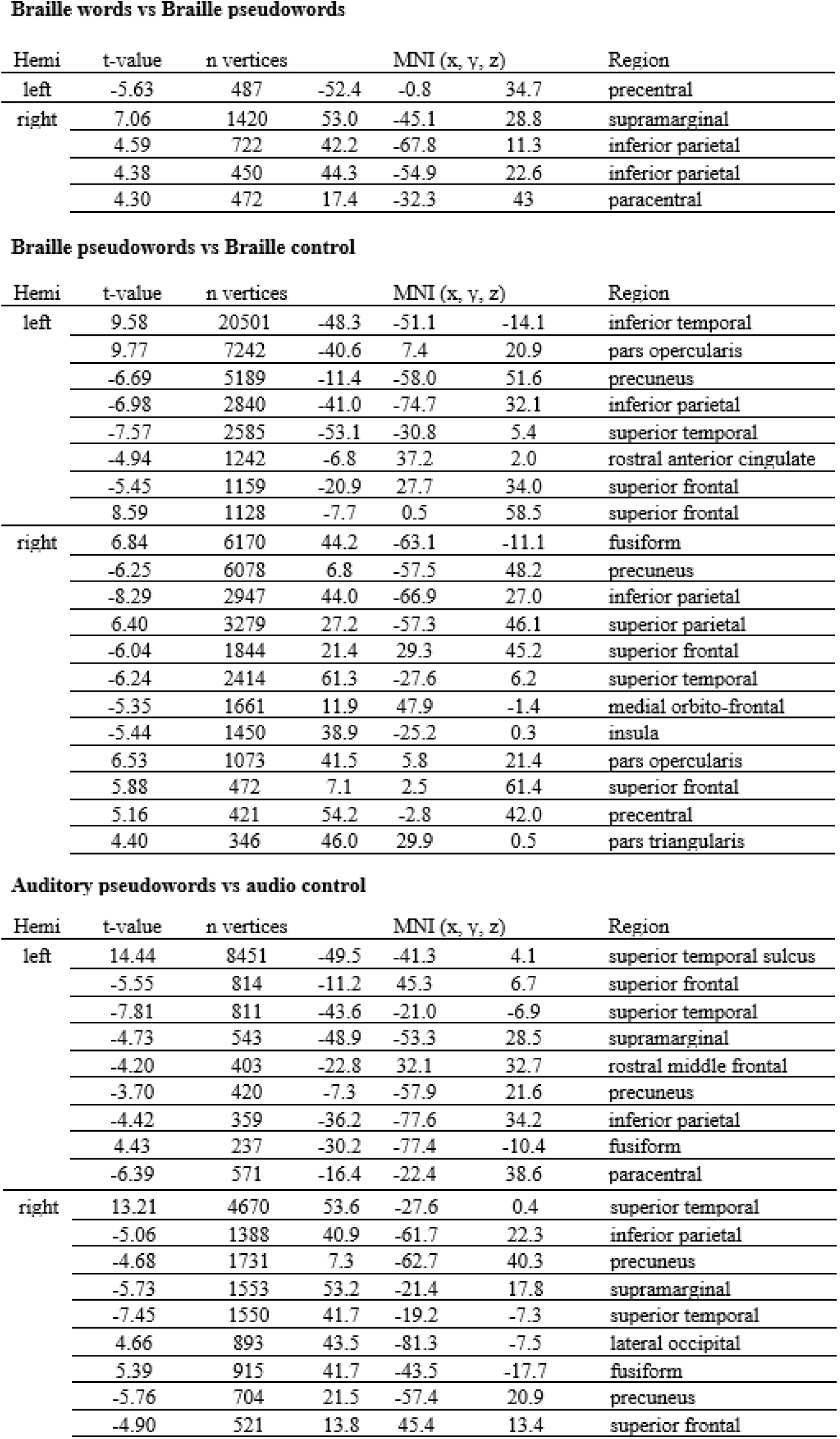
Statistical results of a group analysis (n=38, cluster-forming threshold of z=3 (corresponding to p < 0.001)), corrected for multiple comparisons at p<0.05 using Monte Carlo Simulation, number of permutations = 1000)

Braille words vs Braille pseudowords
| Hemi | t-value | n vertices | MNI (x, y, z) |  |  | Region |
| --- | --- | --- | --- | --- | --- | --- |
| left | -5.63 | 487 | -52.4 | -0.8 | 34.7 | precentral |
| right | 7.06 | 1420 | 53.0 | -45.1 | 28.8 | supramarginal |
|  | 4.59 | 722 | 42.2 | -67.8 | 11.3 | inferior parietal |
|  | 4.38 | 450 | 44.3 | -54.9 | 22.6 | inferior parietal |
|  | 4.30 | 472 | 17.4 | -32.3 | 43 | paracentral |

Braille pseudowords vs Braille control
| Hemi | t-value | n vertices | MNI (x, y, z) |  |  | Region |
| --- | --- | --- | --- | --- | --- | --- |
| left | 9.58 | 20501 | -48.3 | -51.1 | -14.1 | inferior temporal |
|  | 9.77 | 7242 | -40.6 | 7.4 | 20.9 | pars opercularis |
|  | -6.69 | 5189 | -11.4 | -58.0 | 51.6 | precuneus |
|  | -6.98 | 2840 | -41.0 | -74.7 | 32.1 | inferior parietal |
|  | -7.57 | 2585 | -53.1 | -30.8 | 5.4 | superior temporal |
|  | -4.94 | 1242 | -6.8 | 37.2 | 2.0 | rostral anterior cingulate |
|  | -5.45 | 1159 | -20.9 | 27.7 | 34.0 | superior frontal |
|  | 8.59 | 1128 | -7.7 | 0.5 | 58.5 | superior frontal |
| right | 6.84 | 6170 | 44.2 | -63.1 | -11.1 | fusiform |
|  | -6.25 | 6078 | 6.8 | -57.5 | 48.2 | precuneus |
|  | -8.29 | 2947 | 44.0 | -66.9 | 27.0 | inferior parietal |
|  | 6.40 | 3279 | 27.2 | -57.3 | 46.1 | superior parietal |
|  | -6.04 | 1844 | 21.4 | 29.3 | 45.2 | superior frontal |
|  | -6.24 | 2414 | 61.3 | -27.6 | 6.2 | superior temporal |
|  | -5.35 | 1661 | 11.9 | 47.9 | -1.4 | medial orbito-frontal |
|  | -5.44 | 1450 | 38.9 | -25.2 | 0.3 | insula |
|  | 6.53 | 1073 | 41.5 | 5.8 | 21.4 | pars opercularis |
|  | 5.88 | 472 | 7.1 | 2.5 | 61.4 | superior frontal |
|  | 5.16 | 421 | 54.2 | -2.8 | 42.0 | precentral |
|  | 4.40 | 346 | 46.0 | 29.9 | 0.5 | pars triangularis |

| Hemi | t-value | n vertices | MNI (x, y, z) |  |  | Region |
| --- | --- | --- | --- | --- | --- | --- |
| left | 14.44 | 8451 | -49.5 | -41.3 | 4.1 | superior temporal sulcus |
|  | -5.55 | 814 | -11.2 | 45.3 | 6.7 | superior frontal |
|  | -7.81 | 811 | -43.6 | -21.0 | -6.9 | superior temporal |
|  | -4.73 | 543 | -48.9 | -53.3 | 28.5 | supramarginal |
|  | -4.20 | 403 | -22.8 | 32.1 | 32.7 | rostral middle frontal |
|  | -3.70 | 420 | -7.3 | -57.9 | 21.6 | precuneus |
|  | -4.42 | 359 | -36.2 | -77.6 | 34.2 | inferior parietal |
|  | 4.43 | 237 | -30.2 | -77.4 | -10.4 | fusiform |
|  | -6.39 | 571 | -16.4 | -22.4 | 38.6 | paracentral |
| right | 13.21 | 4670 | 53.6 | -27.6 | 0.4 | superior temporal |
|  | -5.06 | 1388 | 40.9 | -61.7 | 22.3 | inferior parietal |
|  | -4.68 | 1731 | 7.3 | -62.7 | 40.3 | precuneus |
|  | -5.73 | 1553 | 53.2 | -21.4 | 17.8 | supramarginal |
|  | -7.45 | 1550 | 41.7 | -19.2 | -7.3 | superior temporal |
|  | 4.66 | 893 | 43.5 | -81.3 | -7.5 | lateral occipital |
|  | 5.39 | 915 | 41.7 | -43.5 | -17.7 | fusiform |
|  | -5.76 | 704 | 21.5 | -57.4 | 20.9 | precuneus |
|  | -4.90 | 521 | 13.8 | 45.4 | 13.4 | superior frontal |

### Dataset 1

This dataset included 25 blind participants (mean age: 36, age SD: 10.04, 16 females; see Table 2) and 25 sighted participants matched for age and sex (mean age: 35.42, age SD: 9.68, 16 females). The data were obtained on a 3T Siemens Trio at the Laboratory of Brain Imaging in the Polish Academy of Sciences in Warsaw. Anatomical images were acquired using a T1-weighted MPRAGE sequence with a 32-channel head coil (176 slices, slice-thickness: 1 mm, TR = 2530 ms, TE = 3.32 ms, flip angle = 7°, matrix size = 256 × 256, voxel size = 1 × 1 × 1 mm). Functional images were acquired using a whole-brain echo-planar imaging sequence with a 12-channel head coil (32 slices, slice-thickness = 4 mm, TR = 2000 ms, TE = 30 ms, flip angle = 80°, matrix size = 64 × 64, voxel size: 3.4 × 3.4 × 4 mm).

### Dataset 2

This dataset included 13 blind participants (mean age: 32.69, age SD: 7.75, 16 females; see Table 2). The data were obtained on the 3T Siemens Magnetom Skyra at the Malopolska Centre of Biotechnology at the Jagiellonian University in Krakow, Poland. Anatomical images were acquired using a T1-weighted MPRAGE sequence with a 32-channel head coil (176 slices, slice-thickness: 0.94 mm, TR = 2300 ms, TE = 2.29 ms, flip angle = 8°, matrix size = 256 × 256, voxel size = 0.94 × 0.9375 × 0.9375 mm). Functional images were acquired using a whole-brain echo-planar imaging sequence with a 32-channel head coil (32 slices, slice-thickness = 4 mm, TR = 2000 ms, TE = 30 ms, flip angle = 80°, matrix size = 64 × 64, voxel size: 3.4 × 3.4 × 4 mm).

### Task and stimuli

During the MRI session, participants underwent structural scanning and performed a language task (Chyl et al., 2021; Dzięgiel-Fivet et al., 2021; Malins et al., 2016). In this task, blind subjects were presented with auditory and tactile stimuli and sighted participants were presented with auditory and visual stimuli. Stimuli in all modalities were presented in three conditions – real words, pseudowords, and a non-linguistic control (see Figure 1). Blind participants were instructed to actively read tactile stimuli whereas sighted participants read the corresponding visual stimuli. Both groups of participants were instructed to actively listen to the auditory stimuli.

**Fig. 1.**
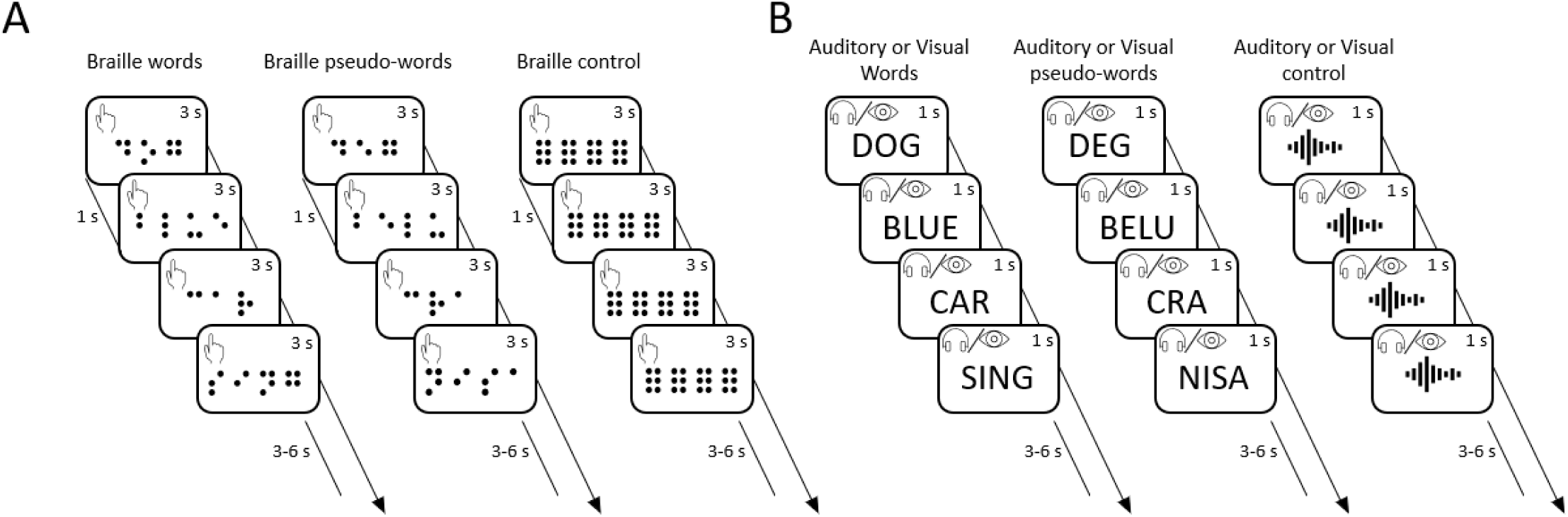
Experimental design. For clarity, stimuli are depicted in English (though they were presented in Polish). (A) Three types of tactile Braille stimuli were presented: words, pseudowords, and a tactile control condition (six dots, no meaning). A block contained 4 stimuli, each presented for 3 seconds with 1 second inter-stimulus interval. Stimuli were presented via a tactile Braille display. (B) Three types of stimuli were presented visually and auditorily: words, pseudowords, and control condition (vocoded words or hashtags). A block contained 4 stimuli, each presented for 1 second without an additional inter-stimulus interval. Stimuli were presented via headphones.

Real words included 148 items which were divided between conditions (i.e., reading and speech comprehension). The conditions were matched in the number of adjectives, verbs and nouns. The words were all of similar word frequency (Fiebach et al., 2002). All items were short (3-4 letters) and consisted of one or two syllables. 148 Pronounceable pseudowords were created to be as similar as possible to the real words, by transposing or substituting letters. All auditory stimuli (including the non-linguistic control) were normalized to last 1000ms. The non-linguistic tactile control stimuli consisted of 3-4 nonsense Braille symbols (all six dots raised), the corresponding visual stimuli consisted of hashtag signs in order to match the Braille condition. The auditory non-linguistic control stimuli consisted of 1 or 2-syllable words (in order to match real and pseudowords length) that were vocoded using Praat (www.praat.org). This process divides the speech signal into three frequency bands, applies the dynamic amplitude contour of the original to a noise source, then recombines these into a unitary signal again. As a result, the auditory stimulus maintains the original dynamic frequency and amplitude pattern, while the phonetic content is significantly diminished.

Each participant completed 3 runs of the task. Each run consisted of 36 blocks. Half of the blocks (18) required listening to auditory stimuli and were 7s long. The other half (18 blocks) required reading. Reading blocks for the blind group required reading braille and were 12s long whereas the sighted control group was reading visual stimuli and each block in this condition lasted 4s. Blocks were evenly distributed across the three conditions (real words, pseudowords and control condition). Within each block, four stimuli from the same condition were presented consecutively. The auditory and visual blocks included 4 trials, each presented for 1000ms with 1000ms interstimulus intervals. The braille blocks included 4 trials, each presented for 3000 ms, with a 1000 ms interstimulus interval. The blocks were separated by breaks, during which a fixation cross was displayed on the screen, lasting between 3000 to 6000 ms (mean = 4000 ms).

The task was programmed using Presentation software (Neurobehavioral Systems, Albany, CA). Auditory stimuli were presented via noise-attenuating headphones (NordicNeuroLab). Tactile stimuli were presented via a NeuroDevice Tacti TM Braille display (Debowska et al., 2013).

### Data analysis

#### Preprocessing

The neuroimaging data were preprocessed using *fMRIPrep* 21.0.2 (Esteban et al., 2018, 2019), which is based on *Nipype* 1.6.1 (Esteban et al., 2022; Gorgolewski et al., 2018).

#### Anatomical data preprocessing

The T1-weighted (T1w) image was corrected for intensity non-uniformity (N4BiasFieldCorrection, (Tustison et al., 2010), distributed (ANTs 2.3.3, (Avants et al., 2008) and used as reference throughout the workflow. The T1w-reference was then skull-stripped (*Nipype* implementation of the antsBrainExtraction.sh workflow from ANTs, using OASIS30ANTs as target template). Brain tissue segmentation was performed on the brain-extracted T1w (fast, FSL 6.0.5.1:57b01774, (Zhang et al., 2001). Brain surfaces were reconstructed (recon-all, FreeSurfer 6.0.1, (Dale et al., 1999) and the previously estimated brain mask was refined (custom variation of the method to reconcile ANTs-derived and FreeSurfer-derived segmentations of the cortical gray-matter of (Klein et al., 2017)). Volume-based spatial normalization to one standard space (MNI152NLin2009cAsym) was performed through nonlinear registration (antsRegistration, ANTs 2.3.3), using brain-extracted versions of both T1w reference and the T1w template. The following template was selected for spatial normalization: *ICBM 152 Nonlinear Asymmetrical template version 2009c* (Fonov et al., 2009).

#### Functional data preprocessing

First, a reference volume and its skull-stripped version were generated (custom methodology of *fMRIPrep*). Head-motion parameters with respect to the blood-oxygen-level-dependent (BOLD) signal reference were estimated (mcflirt, FSL 6.0.5.1:57b01774, (Jenkinson et al., 2002)). BOLD runs were slice-time corrected to 0.971s, 0.5 of slice acquisition range 0s-1.94s (3dTshift from AFNI, (Cox & Hyde, 1997)). The BOLD time-series were resampled onto their original, native space by applying the transforms to correct for head-motion. The BOLD reference was then co-registered to the T1w reference (bbregister, FreeSurfer) which implements boundary-based registration (Greve & Fischl, 2009). Co-registration was configured with six degrees of freedom. Three confounding time-series were calculated based on the *preprocessed BOLD* and were extracted within the CSF, the WM, and the whole-brain masks. Additionally, a set of physiological regressors were extracted to allow for component-based noise correction (*CompCor*, (Behzadi et al., 2007)). Principal components are estimated after high-pass filtering the *preprocessed BOLD* time-series (discrete cosine filter with 128s cut-off). The BOLD time-series were resampled into standard space, generating a *preprocessed BOLD run in MNI152NLin2009cAsym space*. First, a reference volume and its skull-stripped version were generated using a custom methodology of *fMRIPrep*. The BOLD time-series were resampled onto the following surfaces (FreeSurfer reconstruction nomenclature): *fsnative*, *fsaverage*. Gridded (volumetric) resamplings were performed (antsApplyTransforms, ANTs), configured with Lanczos interpolation to minimize the smoothing effects of other kernels (Lanczos, 1964). Non-gridded (surface) resamplings were performed (mri_vol2surf, FreeSurfer). For more details of the pipeline, see the section corresponding to workflows in *fMRIPrep*’s documentation.

#### Univariate GLM Analysis

Prior to first-level modelling, functional data preprocessed with fMRIprep were smoothed with a 5 mm Gaussian kernel. Following that, the smoothed data were submitted for participant-level vertex-wise analysis of blood oxygen level–dependent (BOLD) activation in a general linear model (GLM) using the FreeSurfer Functional Analysis Stream (FS-FAST, FreeSurfer v.7.3.2). A separate GLM was fitted to the left and right hemisphere data, using ordinary least squares estimation. Task-evoked hemodynamic responses associated with the six experimental conditions (i.e. auditory: words, pseudowords, vocoded speech, and: Braille: words, pseudowords, tactile control, or visual: words, pseudowords, visual control) were modeled using a boxcar function convolved with a canonical SPM hemodynamic response function with 0 derivatives. The GLM included one regressor per condition as events of interest. In addition to the experimental condition effects, the GLM design included second order polynomial regressors to remove slow trends as well as 6 subject-specific motion parameters estimated by fMRIprep. To identify the effects of interest for our study, we modeled the following first-level contrasts. First, to identify semantic processing on the participant level, we contrasted the processing of words with the processing of pseudowords in both modalities. Second, to identify phonological processing, we contrasted the processing of pseudowords with the processing of the control condition in the auditory modality. Finally, contrasting pseudowords with the control condition in Braille reading in the blind group and in visual reading in the sighted group, we identified the orthographic processing. Contrasts on the individual level were then submitted to the second-level analysis together with a covariate of no interest of the scanning site to account for potential variance associated with differences in equipment used and, possibly, experimenter bias. The second-level statistical maps for each contrast were corrected for multiple comparisons using a non-parametric cluster-wise correction across two spaces (left and right hemisphere) using permutation testing (1000 permutations, cluster forming threshold > 3, p<0.05).

#### Analysis of associations between cortical thickness and functional activations

Individual functional activation maps and cortical thickness maps were resampled and mapped to the FreeSurfer surface template (fsaverage) and concatenated into a single file per hemisphere for statistical analysis. Vertex-wise cortical thickness measurements were extracted from each participant’s occipital cortex. The reconstructed cortical surface was automatically parcellated for each participant into the 74 neocortical areas in each hemisphere defined by the Destrieux atlas (Destrieux et al., 2010). A total of 15 areas were defined as anatomical regions of interest (ROI) in each hemisphere: the occipital pole, the inferior occipital gyrus and sulcus, the cuneus, the fusiform gyrus, the anterior occipital sulcus, the lingual gyrus, the middle occipital gyrus, the superior occipital gyrus, the calcarine sulcus, the posterior transverse collateral sulcus, the lateral occipito-temporal sulcus, the collateral and lingual sulci, the middle occipital sulcus and the lunatus sulcus, the superior occipital and transverse occipital sulci, as well as the fissure (see Figure 2). Then, for each of the contrasts, percent signal change was extracted for each vertex included in the ROIs. As a result, for each of the 38 blind participants we had one measure of cortical thickness and four measures of percent signal change (one for each contrast) for each vertex.

**Figure 2.**
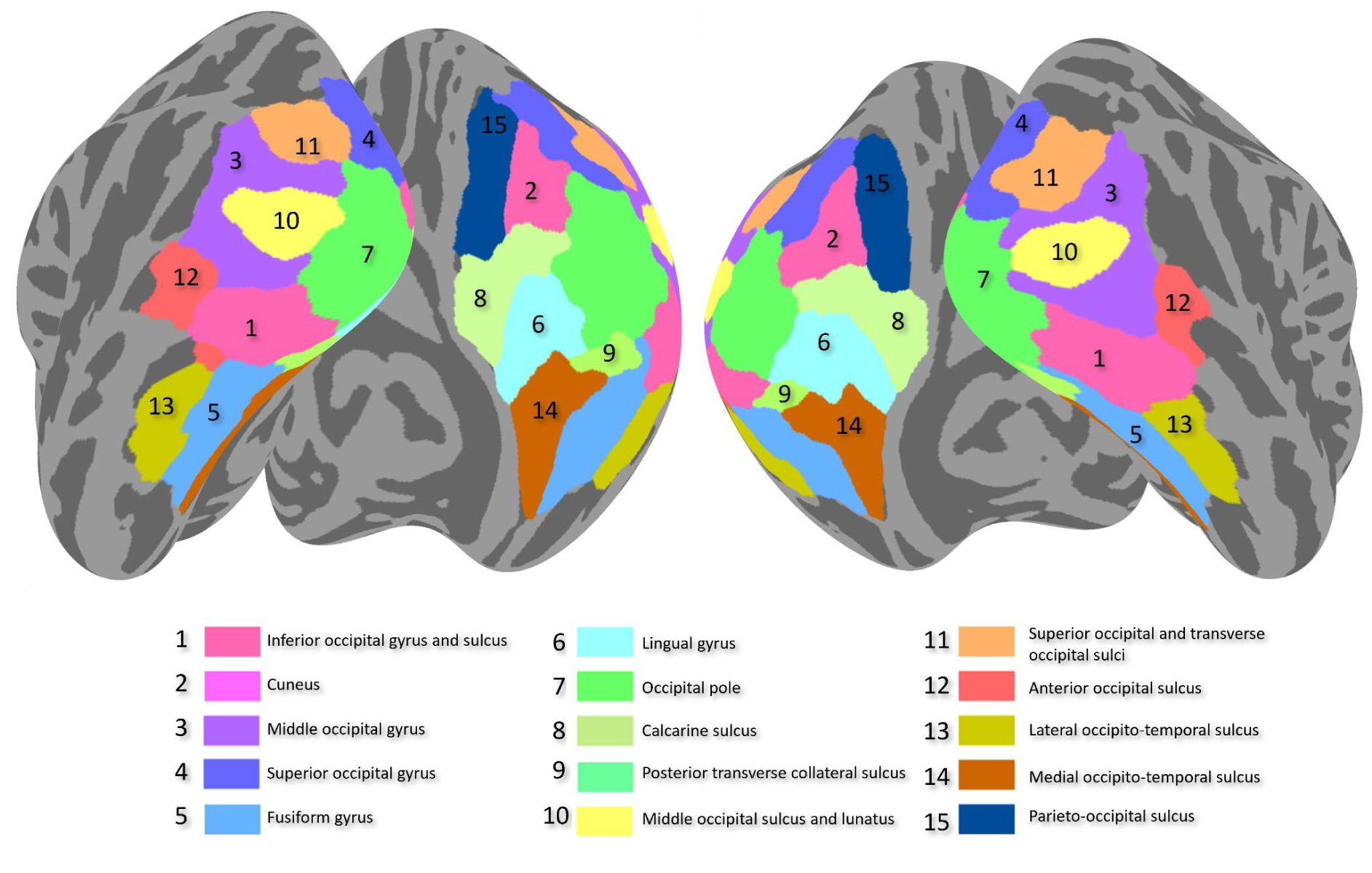
Regions of interest on the medial and lateral surface of the occipital cortex based on the parcellations of the Destrieux atlas (Destrieux et al., 2010).

**Figure 3.**
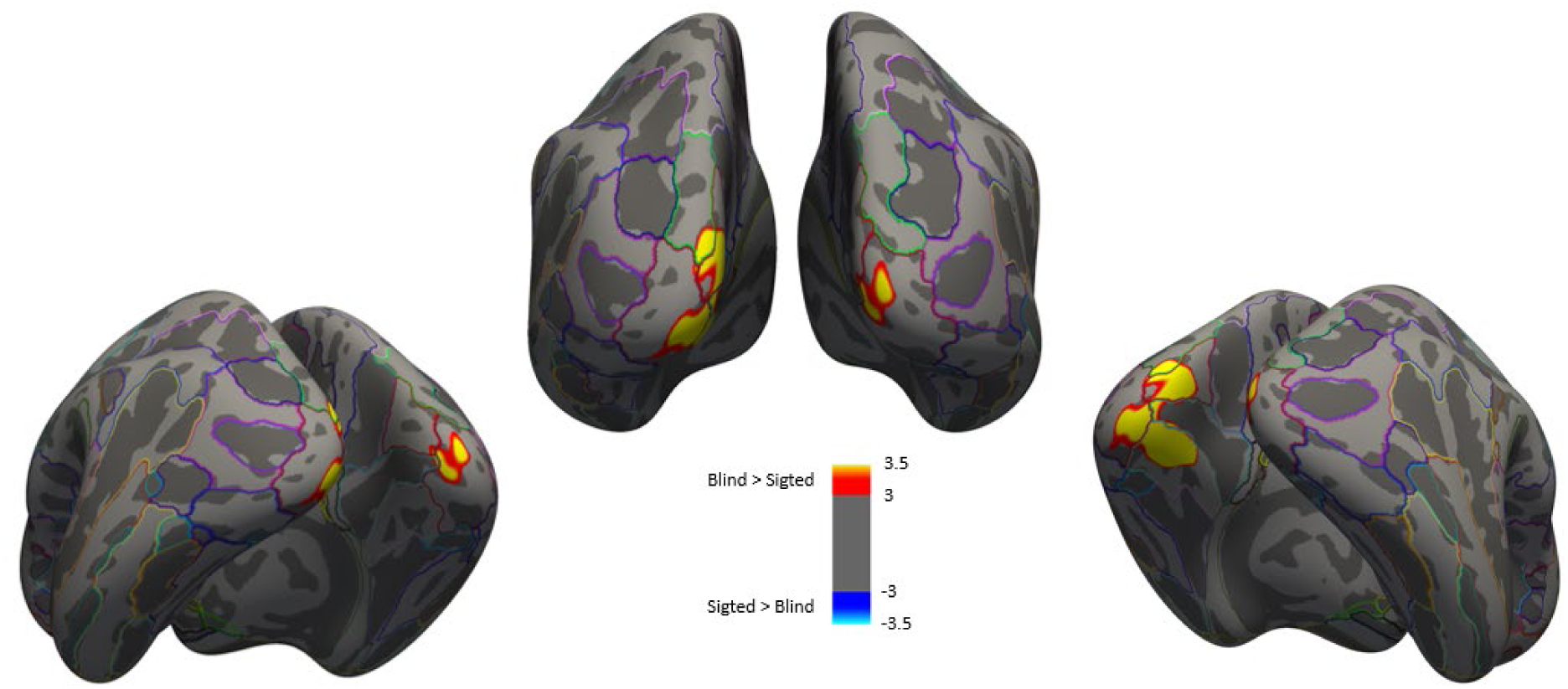
Statistical maps of brain regions that differ in cortical thickness between the blind group (n=25) and the sighted controls (n=25). Results are displayed at a cluster-forming threshold of z=3 (corresponding to p<0.001), corrected for multiple comparisons at p<0.05 using Monte Carlo Simulation, number of permutations = 1000). Color scale codes -log10(p). Maps are presented on the inflated surface (dark gray: sulci, light gray: gyri) of the FreeSurfer standard brain.

Statistical analyses were performed in R (R Core Team, 2021) using the R nmle package (v:3.1-162, (Pinheiro & Bates, 2000)). Separate linear mixed models were run for each ROI and contrast combination, resulting in 60 models. For model selection we followed a protocol described in (Zuur et al., 2009). Percent signal change was modeled as a continuous response variable in a linear mixed-effects model at vertex resolution. The feasibility of applying a single model to the data from all 15 ROIs was dismissed due to the implications of incorporating a 15-level factor, potentially involving interactions, in both the fixed and random components of the model, as this would have resulted in an exponential increase in the number of model parameters. Instead, the model was fitted separately to the data of every combination of contrast and ROI. The contrast- and ROI-specific model included fixed effects for cortical thickness, hemisphere, age, sex, and scanner, and two interaction terms, one for cortical thickness and hemisphere, and one for age and sex. The random part contained two levels of nested random effects. The outer level for the subject factor contained a random intercept. The inner level for the hemisphere factor was nested within subject and contained a random intercept plus a random slope for cortical thickness which were assumed uncorrelated with equal variances in both hemispheres. The variance of the within-group errors was assumed to be different for scanner 1 and 2. P-values were adjusted using per-contrast Bonferroni corrections.

In the initial data collection (Warsaw scanning site), fMRI data were also collected from a group of sighted controls. This data was analyzed to assess whether possible associations were specific to the blind group. For this comparison, we analyzed only a subset of the data from blind participants for whom we had corresponding control matches (n = 25). A similar analysis to that described above was conducted on this subset. However, in this case, an additional variable for group (blind vs sighted control) as well as its interactions with hemisphere and cortical thickness were included in the LMM, while the scanner variable was dropped.

## RESULTS

### Difference in cortical thickness between early blind and sighted participants

We examined cortical thickness differences between early blind individuals and sighted controls (Figure 2). A bilateral cluster in the early visual cortex showed significantly greater thickness in the early blind (cluster-forming threshold of z=3, corrected for multiple comparisons at p<0.05 using Monte Carlo simulation). In the left hemisphere, the cluster extended from the cuneus through the occipital pole to the lingual gyrus. In the right hemisphere, a homologous but smaller cluster was found in the occipital pole. The reverse contrast (sighted controls > early blind) did not yield significant results. The regions showing thickness differences align well with previous voxel-based morphometry studies in the early blind (Jiang et al., 2009; Noppeney et al., 2005; Park et al., 2009; Ptito et al., 2008).

### Functional MRI data analysis in the group of early blind participants

Whole-brain analysis revealed activations evoked by different levels of linguistic processing. Activation evoked by semantic processing was assessed separately in Braille and auditory stimuli by contrasting words against pseudowords. In Braille reading this contrast revealed activation primarily in right parietal regions, including the superior temporal sulcus and the angular gyrus but also involved areas in the right middle occipital gyrus (see Figure 4A and Table 3). The opposite contrast (shown in blue) revealed increased activation in left precentral motor areas, possibly reflecting processes related to covert articulation. Contrasting words against pseudowords in the auditory modality did not result in any significant activations (see Figure 4C). Additional contrasts are presented in the supplementary materials (see Supplementary Figure 2).

**Figure 4.**
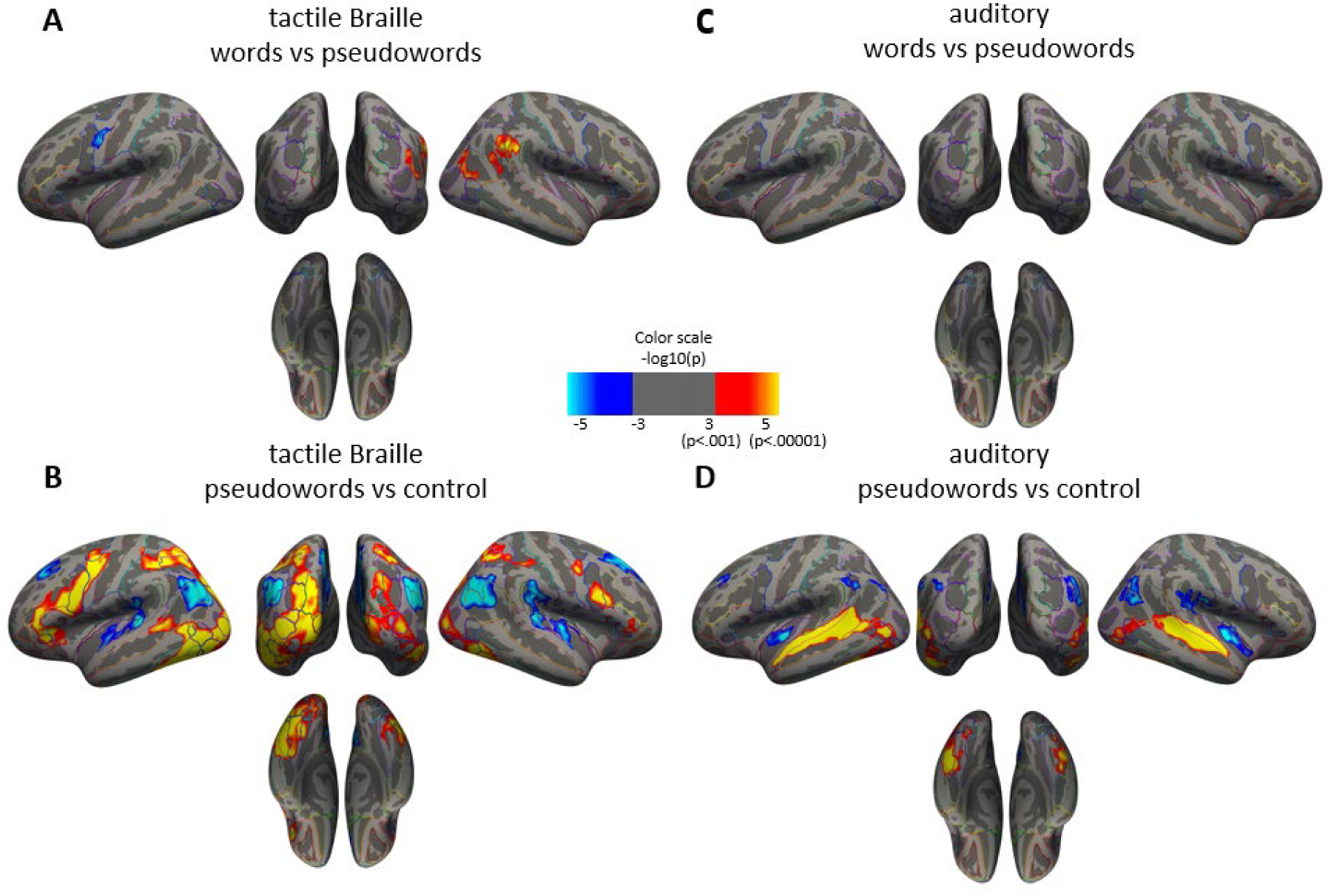
Statistical maps of brain activations related to **A**) semantic processing in Braille (contrast: Braille words vs Braille pseudowords), **B**) orthographic processing in Braille (contrast: Braille pseudowords vs Braille control), **C**) semantic processing in spoken words (contrast: audio words vs audio pseudowords), and **D**) Phonological processing in spoken words (contrast: audio pseudowords vs audio control). Color scale codes -log10(p). Maps are superimposed on the inflated surface (dark gray: sulci, light gray: gyri) of the FreeSurfer standard brain, displayed at a cluster-forming threshold of z=3 (corresponding to p < 0.001), corrected at p<0.05 for multiple comparisons using Monte Carlo Simulation, number of permutations = 1000).

**Figure 5.**
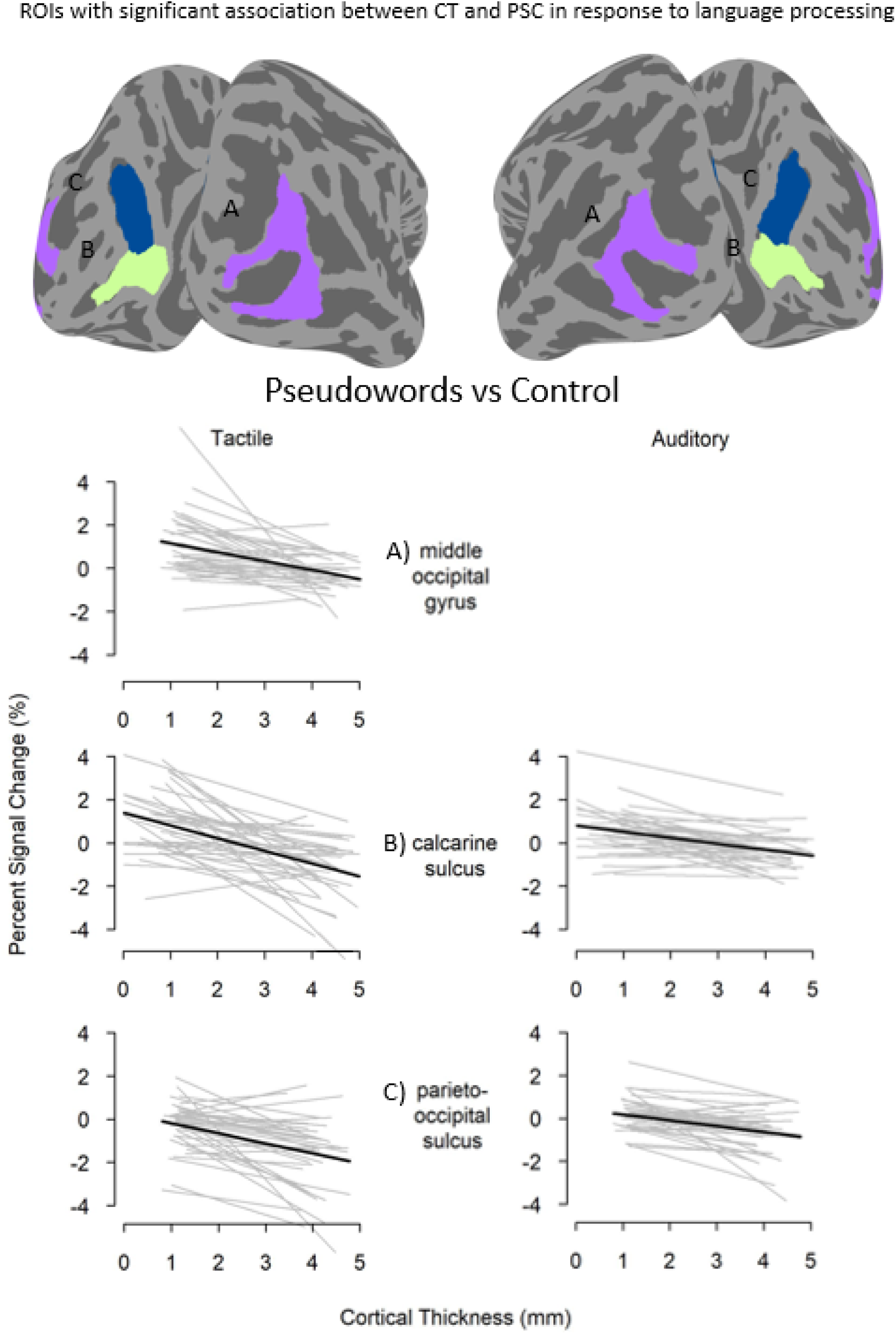
Top panel: Cortical regions with significant association between cortical thickness and functional activation in the early blind group (n=38). The bottom plots show fitted curves of contrast- and ROI-specific linear mixed-effects models in which the main effect for cortical thickness was significant at the 5% level after a per-contrast Bonferroni correction. Curves represent the relationship between cortical thickness and percent signal change. Within-group fitted curves for left and right hemispheres nested within-subject were averaged to produce one gray line per subject. The thick, black line represents the fitted values for the population. Every combination of hemisphere, age, sex, and scanner results in a different intercept, shifting the population curve up or down. To visually center the black line with respect to the gray lines the average intercept computed over subjects was used.

**Table 3.**
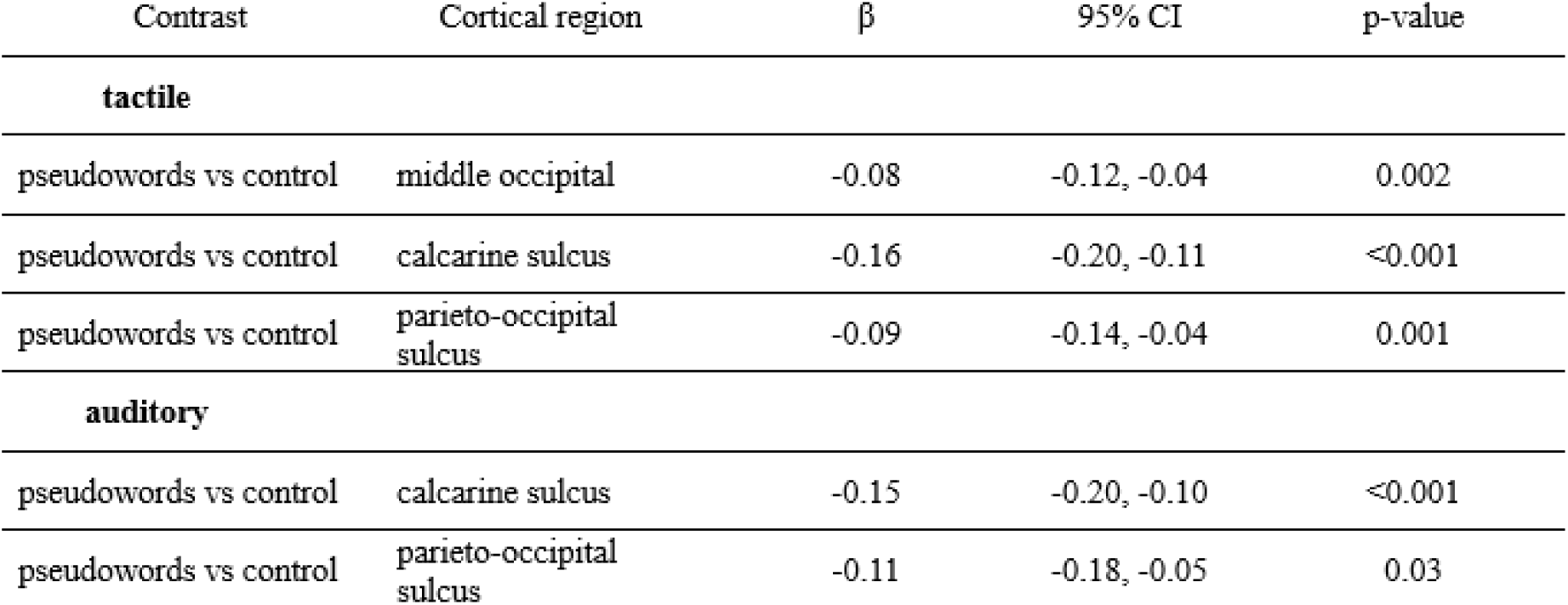
Cortical regions showing a significant association between cortical thickness and percent signal change in the early blind group (n=38). P-values were adjusted using a per-contrast Bonferroni correction.

| Contrast | Cortical region | $\beta$ | 95% CI | p-value |
| --- | --- | --- | --- | --- |
| **tactile** |  |  |  |  |
| pseudowords vs control | middle occipital | -0.08 | -0.12, -0.04 | 0.002 |
| pseudowords vs control | calcarine sulcus | -0.16 | -0.20, -0.11 | <0.001 |
| pseudowords vs control | parieto-occipital sulcus | -0.09 | -0.14, -0.04 | 0.001 |
| **auditory** |  |  |  |  |
| pseudowords vs control | calcarine sulcus | -0.15 | -0.20, -0.10 | <0.001 |
| pseudowords vs control | parieto-occipital sulcus | -0.11 | -0.18, -0.05 | 0.03 |

To assess orthographic processing during Braille reading, we contrasted pseudowords with the control condition. This revealed bilateral clusters of activation in the postcentral and the inferior frontal sulcus, bilateral clusters in the superior parietal lobule extending into the intraparietal sulcus and occipito-temporal regions, including the cuneus, the occipital pole, and the fusiform gyrus, as well as bilateral clusters in the superior temporal sulcus, and the insular cortex. Clusters of activation were more extensive in the left than in the right hemisphere (see Figure 4B and Table 3). Additionally, we observed activation in the opposite contrast (shown in blue) in the frontal cortex, perisylvian areas, and the angular gyrus. We also identified a cluster in the medial plane, encompassing the parieto-occipital sulcus, precuneus, and subparietal sulcus (see Figure 4B and Table 3).

Phonological processing (pseudowords vs control in the auditory modality) evoked increased activation within the superior temporal cortex, the anterior and inferior occipital sulci, and the inferior occipital gyrus extending into the lateral occipito-temporal sulcus and the fusiform gyrus. This cluster appeared bilaterally, although more extensively on the left side. Additionally, we observed a cluster of activation that included the occipital pole, the calcarine sulcus, and the cuneus in the left hemisphere. We observed activations in response for the opposite contrast (shown in blue) within the superior and middle frontal gyri and a cluster in the right angular gyrus, the superior temporal sulcus, and the middle occipital gyrus (see Figure 2, and Table 3).

See additional analyzes in the supplementary materials (Supplementary Figure 2).

### Negative Association Between Cortical Thickness and Activation in the group of early blind participants

The relationships between cortical thickness and percent signal change in the early blind group were tested separately for each ROI and contrast. We found negative associations between cortical thickness and percent signal change in the occipital cortex of blind participants. For phonological processing in the auditory modality, lower cortical thickness was related to increased functional activation in the calcarine sulcus and in the parieto-occipital sulcus (see Figure 4 and Table 4). For orthographic processing in the tactile modality, this effect was significant in the middle occipital gyrus, the calcarine sulcus, and the parieto-occipital sulcus (see Figure 4 and Table 4). We did not find significant associations between the cortical thickness and the percent signal change in the semantic processing (words vs pseudowords).

## Discussion

Previous studies have reported that the structure and function of the occipital cortex is drastically altered in blind individuals. However, how these changes in structure and function relate to each other has remained unclear. Here, we investigated the association between cortical thickness and the functional activation in the occipital cortex of blind individuals during linguistic processing. Consistent with our predictions and previous findings (Anurova et al., 2015), we observed negative associations between cortical thickness and the functional response to various levels of language processing in the occipital cortex of blind, but not sighted individuals. That is, the thinner the occipital cortex, the greater the activation in response to linguistic information in blind individuals. Thus, our study provides unique insights into theoretical debates on the apparent increased cortical thickness in blind individuals.

### Cross-modal plasticity

By now, a plethora of studies have reported that the occipital cortices, which are typically associated with visual processing in sighted individuals, are activated during auditory and tactile processing in blind individuals(Bedny et al., 2011; Collignon et al., 2011; Dzięgiel-Fivet et al., 2021; Röder et al., 2002; Sadato et al., 1996). Consistent with previous studies, we observed that auditory and tactile linguistic processing activated the occipital cortex of early blind individuals. We found increased activation in response to semantic processing during Braille reading in parietal regions in the right hemisphere, including the inferior parietal lobule, the supramarginal gyrus, and the posterior parts of the cingulate gyrus (see Figure 4A and Table 3). We did not observe activations in classical language areas that are typically activated during semantic processing in sighted individuals (Binder et al., 2009), such as the superior and middle temporal cortex. This is consistent with previous studies that have investigated various levels of linguistic processing in blind individuals and have reported little or reduced activation in classic language regions in blind compared to sighted individuals (Burton et al., 2002; Dzięgiel-Fivet et al., 2021; Gizewski et al., 2003; Kim et al., 2017). This result could suggest that the linguistic functions of the temporal cortex in blind individuals during reading may, at least partially, be taken over by other regions, including the occipital cortex.

Overall, we observed weak activations when we contrasted the reading of Braille words with the reading of Braille pseudowords. No significant activation for this contrast was observed in the left vOTC. It has been argued that the vOTC activation is greater for pseudowords than for words because of increased prediction error (Price & Devlin, 2011). The opposite contrast (Pseudowords contrasted with words) elicited increased activations in the inferior frontal cortex and the precentral gyrus (Supplementary Figure 1). It has been suggested that increased activation for pseudowords vs words in this area reflects the increased demands on sublexical conversion of orthography to phonology (Hagoort et al., 1999). Alternatively, it has been suggested that these effects are related to general task difficulty and are not specific to linguistic processing (Fedorenko et al., 2013). This was supported by Fiez and Petersen (1998) as well as Fiez et al. (1999), who observed increased activation in the left inferior frontal cortex when comparing the reading of words with irregular spellings (e.g., KNIFE) to those with regular spellings (e.g., BROOM) (Fiez et al., 1999; Fiez & Petersen, 1998). Increased activation in the left inferior frontal cortex for pseudowords vs words has also been observed when the words are presented auditorily (Xiao et al., 2005), corroborating the notion that this effect reflects task difficulty and not necessarily grapheme-to-phoneme conversion.

When we compared the processing of pseudowords to the processing of the non-linguistic control items in Braille reading, we observed increased activation within the classic language network, including the postcentral sulcus and perisylvian areas. This result is consistent with studies in sighted individuals (Schuster et al., 2015). In addition, we observed increased activation in parietal and occipito-temporal regions. The same contrast (pseudowords vs control) in the auditory modality in the group of blind participants revealed activation in the classic perisylvian language areas, as well as in occipital areas. These findings provide further evidence for the notion that at least some linguistic processes, possibly phonological or orthographic, are taken over by the occipital areas in blind individuals. However, we cannot determine the specific type of linguistic processing involved. It is important to note that silent reading of pseudowords, as pronounceable strings, could also involve phonological processing by the process of grapheme to phoneme conversion (Clifton, 2015). Therefore, our design does not allow us to fully distinguish between orthographic and phonological aspects of reading.

The results of this study should be considered in the context of the differences between Braille reading in blind individuals and print reading in sighted individuals. These processes are different on many levels starting from the low-level processing up to social aspects. Braille reading is slower, sequential, and requires the volitional aspect of moving the finger across the text. Braille reading speed in Polish language has been reported to be 60.5 words per minute (Rączy et al., 2019). Conversely, in sighted individuals, print reading of words up to 6 letters seems to be automatic and fast (Cohen et al., 2008). The average print reading speed has been reported to be 242 words per minute (Brysbaert, 2019), making it roughly four times as fast as Braille reading. It should also be kept in mind that currently blind individuals often prefer other forms of communication that are more easily accessible, such as audio recordings, audio books, and/or audio descriptions. As a result, blind individuals tend to read less frequently than sighted individuals which could result in neural differences between the two groups.

### Relationship between structure and function

We found that thinner occipital cortices in blind individuals were associated with increased functional activation during various aspects of linguistic processing. This is in line with previous studies that have reported a similar association in the occipital cortex of blind individuals who performed low-level auditory tasks (Anurova et al., 2015) and fronto-parietal areas of children who performed linguistic tasks (Lu et al., 2009; Nuñez et al., 2011). However, while Anurova et al. (2015) focused on low-level auditory tasks, our study extends this discussion by demonstrating similar associations in high-level language tasks. Moreover, in our analysis we tested the associations between structure and function at the single-subject level by fitting the model to vertex-wise cortical thickness and percent signal change within each region of interest. Thus, our results provide new insights into the role of the occipital cortex in language processing in early blind persons even at the single subject level.

The results from additional analysis performed on a subset of the blind participants were consistent with the findings from the full group analysis. Three out of five contrast-ROI combinations from the initial analysis showed significant differences between the groups. In those cases, the associations are negative in the blind group, as predicted, and non-significant in the control group (Supplementary Table 1 and Supplementary Figure 3). Two additional contrast-ROI combinations show significant group differences where the association between the cortical thickness and the percent signal change is not significant in the blind group but is positive in the control group. These results align with our initial findings, and the differences are likely reflective of the reduced sample size. These findings suggest that the relationship between cortical thickness and functional activation differs between sighted and blind individuals.

### Possible mechanisms

Cross-sectional histological studies on post-mortem brains have shown that inefficient synapses, dendrites, and neurons are eliminated during early brain development (Huttenlocher et al., 1982; Petanjek et al., 2008; Rakic et al., 1986). It has been suggested that such pruning mechanisms play a crucial role in functional maturation by optimizing synaptic efficiency (Rakic et al., 1986) and may lead to cortical thinning during development (Huttenlocher et al., 1982; Huttenlocher & Dabholkar, 1997) . It has been suggested that the apparent thicker occipital cortices in blind individuals may be a reflection of a disruption of synaptic pruning (Jiang et al., 2009). However, the prevailing view that the cortex undergoes thinning during development has recently been challenged by studies using quantitative MRI and diffusion MRI. Instead, these studies suggest that the cortex becomes more myelinated, affecting the contrast of MR images and, thus, the apparent gray-white matter boundary (Gomez et al., 2017; Natu et al., 2019). Based on these findings it could also be hypothesized that what previous studies have identified as a thicker visual cortex in blind individuals, is actually reflective of a reduction in myelination of these regions in blind individuals.

Developmental studies have shown that a thinner cortex is associated with stronger activations (Lu et al., 2009; Nuñez et al., 2011). In line with this, we observed a negative association between cortical thickness and percent signal change in congenitally blind individuals, that is, thinner occipital cortex was associated with stronger activations. Thus, our data suggests that effective crossmodal recruitment of the occipital cortex may require some degree of pruning. Moreover, it could be hypothesized that a thinner cortex is more myelinated, which in turn may lead to increased crossmodal recruitment of the affected area (Gomez et al., 2017; Natu et al., 2019).

The occipital cortices of blind individuals are recruited in various non-visual processes (Bedny et al., 2011; Dzięgiel-Fivet et al., 2021; Hölig et al., 2014; Sadato et al., 1996). Thus, it seems logical to assume that these brain circuits undergo pruning and/or myelination to optimize synaptic efficiency for these new non-native inputs (Anurova et al., 2015). While our study cannot disentangle between these two mechanisms, it could be hypothesized that one or both of these two developmental mechanisms is necessary for effective functioning of occipital areas in blind individuals. Future studies using a combination of quantitative, functional, and diffusion MRI are necessary to shed light on this issue.

## Conclusion

The present study provides further evidence for functional reorganization of the occipital cortices of early blind individuals. This functional reorganization seems to be related to structural characteristics of the occipital cortex; we observed that thinner occipital cortex goes hand in hand with increased functional activation during linguistic processing in early blind individuals. Furthermore, these results suggest that developmental mechanisms such as pruning and/or myelination play a critical role in the functional maturation of brain circuits.

## Conflicts of interest

The authors declare no conflicts of interest.

## Author Contributions

**MC**: Conceptualization, Methodology, Software, Formal analysis, Investigation, Writing - Original Draft, Writing - Review & Editing,

**ALS:** Conceptualization, Methodology, Formal analysis, Writing - Original Draft, Writing - Review & Editing, Supervision,

**KH:** Conceptualization, Methodology, Formal analysis, Investigation, Writing - Review & Editing,

**AW:** Conceptualization, Methodology, Software, Formal analysis, Writing - Review & Editing,

**CB:** Methodology, Software, Formal analysis, Writing - Review & Editing,

**JB:** Conceptualization, Methodology, Software, Investigation, Writing - Review & Editing,

**JV:** Conceptualization, Methodology, Software, Writing - Review & Editing,

**OC:** Conceptualization, Methodology, Writing - Review & Editing, Supervision,

**KJ:** Conceptualization, Methodology, Resources, Writing - Review & Editing, Supervision, Funding acquisition,

**MS:** Conceptualization, Methodology, Resources, Writing - Review & Editing, Supervision, Funding acquisition

## Acknowledgments

We would like to thank our participants without whom the study would not have been possible. We would like to express our heartfelt gratitude to Gabriela Dzięgiel for her invaluable contributions to the methodology and her dedicated efforts in recruitment and data collection.

## Funding

This work was supported by the Polish National Science Centre (NCN; grant number 2018/30/A/HS6/00595 to M.S.). Clemens Baumbach was supported by the project “NeuroSmog: Determining the impact of air pollution on the developing brain” (Nr. POIR.04.04.00–1763/18–00) which is implemented as part of the TEAM-NET programme of the Foundation for Polish Science, co-financed from EU resources, obtained from the European Regional Development Fund under the Smart Growth Operational Programme.

## Supplementary materials

**Supplementary Figure 1.**
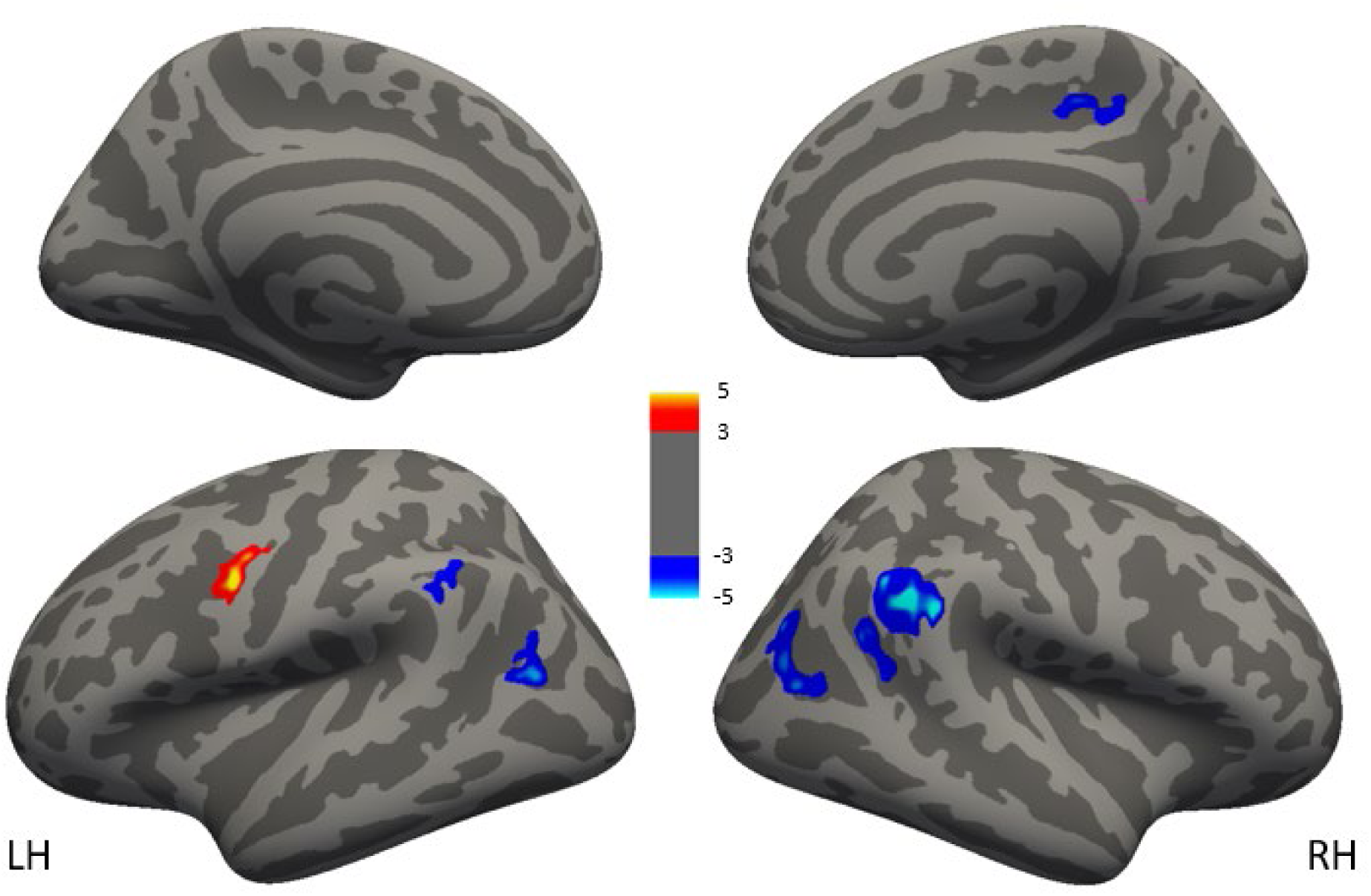
Statistical maps of brain activations evoked by the contrast: Braille pseudowords vs Braille words, Maps are presented on the inflated surface (dark gray: sulci, light gray: gyri) of the FreeSurfer standard brain, displayed at a cluster-forming threshold of z=3 (corresponding to p < 0.001), corrected for multiple comparisons at p<0.05 using Monte Carlo Simulation, number of permutations = 1000.

**Supplementary Figure 2.**
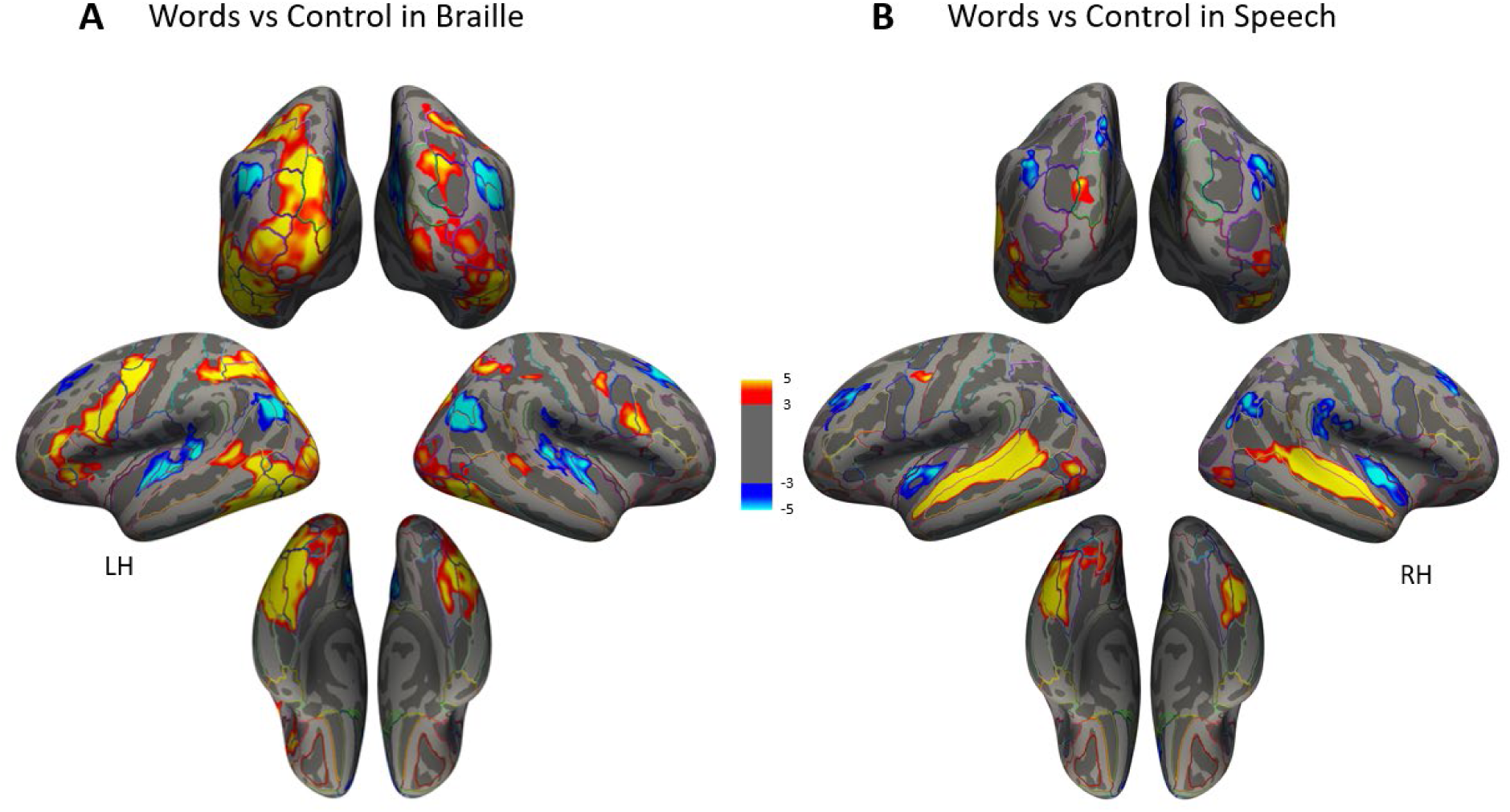
Statistical maps of brain activations evoked by the contrast: words vs control condition in Braille and spoken stimuli in the blind group. Maps are presented on the inflated surface (dark gray: sulci, light gray: gyri) of the FreeSurfer standard brain, displayed at a cluster-forming threshold of z=3 (corresponding to p < 0.001), corrected for multiple comparisons at p<0.05 using Monte Carlo Simulation, number of permutations = 1000.

The Association Between Cortical Thickness and Activation differs Between Blind and Sighted individuals

For orthographic processing (reading pseudowords vs reading control), the group of blind individuals, but not the group of sighted controls, showed a negative relationship between cortical thickness and functional activation in the calcarine sulcus. For phonological processing in the auditory domain (auditory pseudowords vs auditory control), a significant group difference was found in the calcarine sulcus. Here, the blind, but not the sighted group, showed a negative relationship between cortical thickness and functional activation.

**Supplementary Table 1.**
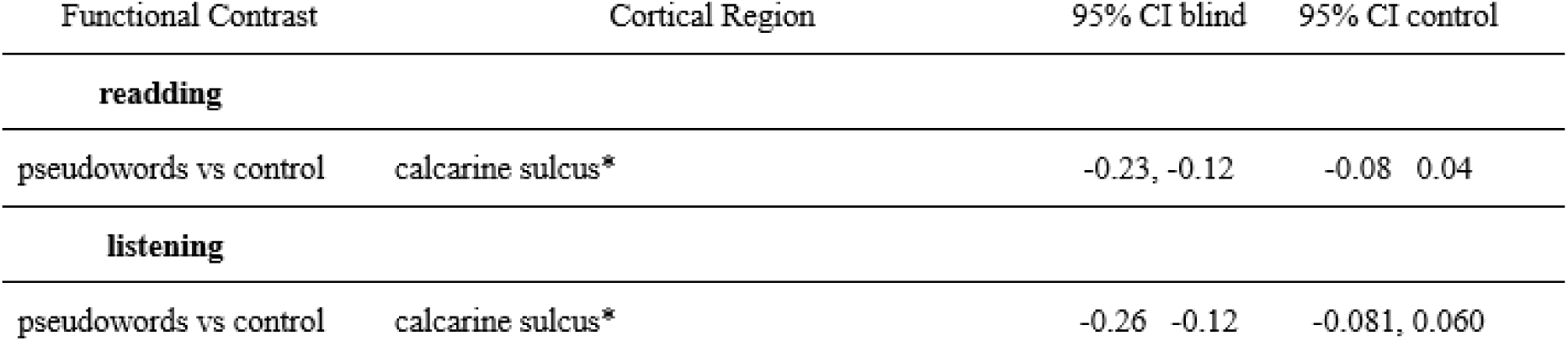
Cortical regions showing a significant difference in the association between cortical thickness and percent signal change between the groups (* marks region-contrast combinations that showed significant associations between the percent signal change and cortical thickness in the previous analysis of the early blind group). The table presents estimates for contrasts in cortical regions which showed a significant interaction between group and cortical thickness. Confidence intervals for the trends as a function of group and contrast.

**Supplementary Figure 3.**
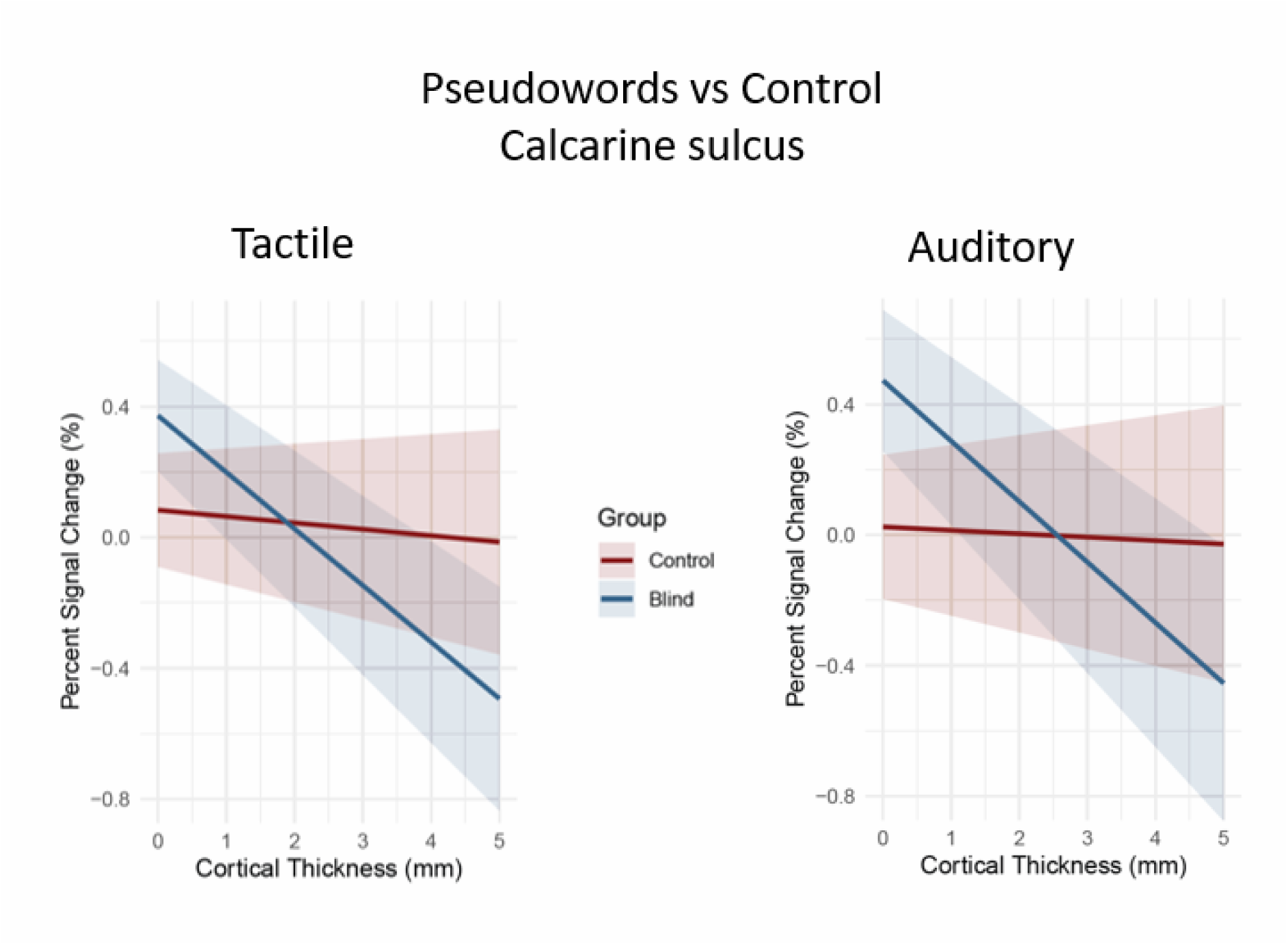
Cortical regions with significant interaction between cortical thickness and group (blind group n=25, control group n=25). The plots show population-averaged fitted curves of contrast- and ROI-specific linear mixed-effects models in which the main effect for cortical thickness was significant at the p<.05 level after a per-contrast Bonferroni correction. The ribbons represent confidence intervals.

## Notes

### Competing Interest Statement

The authors have declared no competing interest.

