## Supplementary material for "Association between Cortical Thickness and Functional Response to Linguistic Processing in the Occipital Cortex of Early Blind Individuals": Suplementary Materials

### Supplementary materials

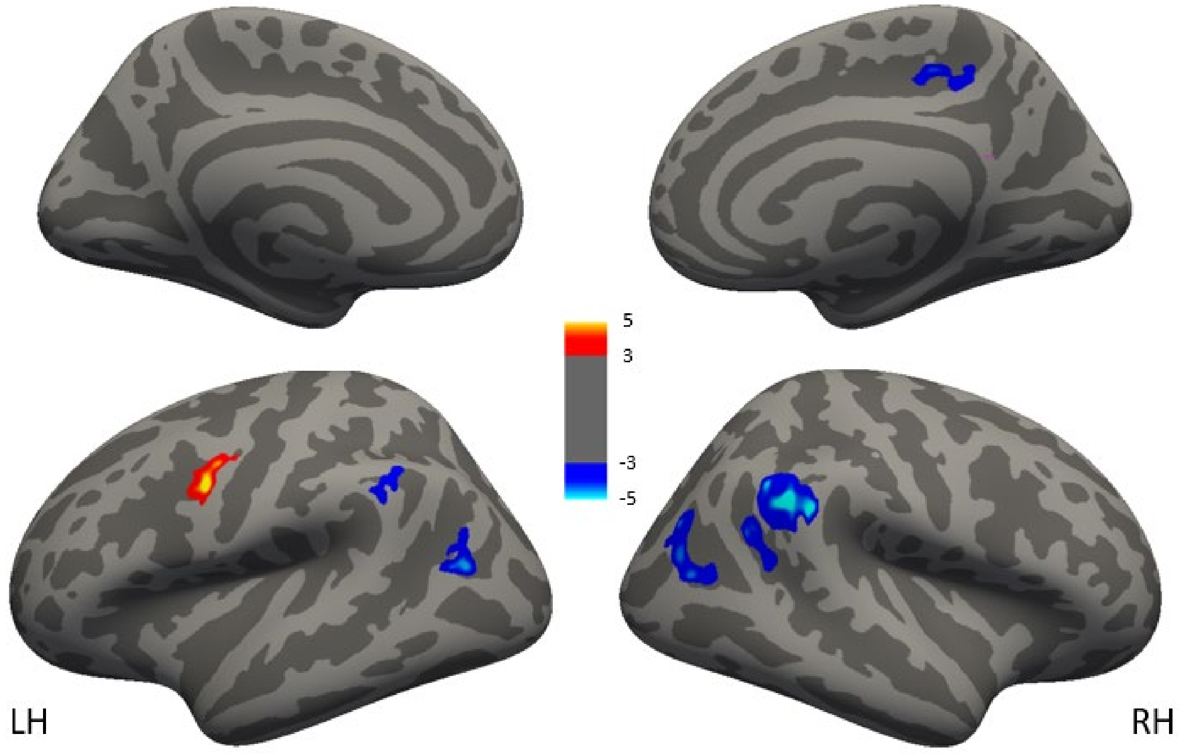


*Supplementary Figure 1. Statistical maps of brain activations evoked by the contrast: Braille pseudowords vs Braille words, Maps are presented on the inflated surface (dark gray: sulci, light gray: gyri) of the FreeSurfer standard brain, displayed at a cluster-forming threshold of z=3 (corresponding to p < 0.001), corrected for multiple comparisons at p<0.05 using Monte Carlo Simulation, number of permutations = 1000.*

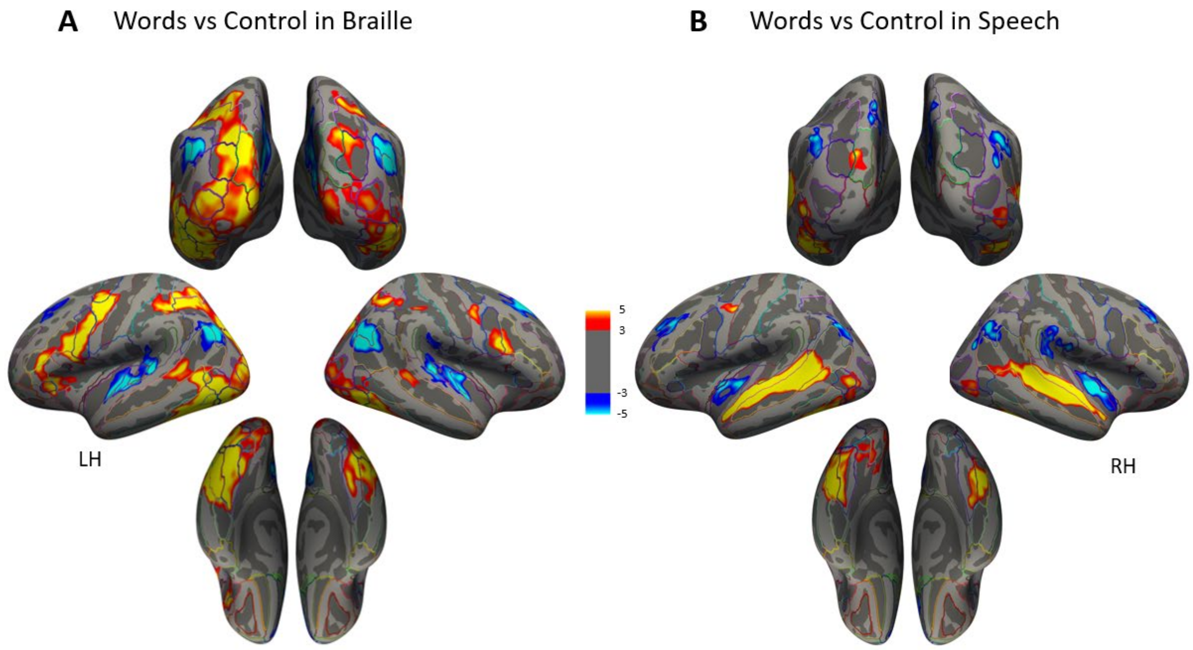


*Supplementary Figure 2. Statistical maps of brain activations evoked by the contrast: words vs control condition in Braille and spoken stimuli in the blind group. Maps are presented on the inflated surface (dark gray: sulci, light gray: gyri) of the FreeSurfer standard brain, displayed at a cluster-forming threshold of z=3 (corresponding to p < 0.001), corrected for multiple comparisons at p<0.05 using Monte Carlo Simulation, number of permutations = 1000.*

#### The Association Between Cortical Thickness and Activation differs Between Blind and Sighted individuals

For orthographic processing (reading pseudowords vs reading control), the group of blind individuals, but not the group of sighted controls, showed a negative relationship between cortical thickness and functional activation in the calcarine sulcus. For phonological processing in the auditory domain (auditory pseudowords vs auditory control), a significant group difference was found in the calcarine sulcus. Here, the blind, but not the sighted group, showed a negative relationship between cortical thickness and functional activation.

| Functional Contrast | Cortical Region | 95% CI blind | 95% CI control |
|---|---|---|---|
| **readding** | | | |
| pseudowords vs control | calcarine sulcus* | -0.23, -0.12 | -0.08 0.04 |
| **listening** | | | |
| pseudowords vs control | calcarine sulcus* | -0.26 -0.12 | -0.081, 0.060 |

*Supplementary Table 1. Cortical regions showing a significant difference in the association between cortical thickness and percent signal change between the groups (* marks region-contrast combinations that showed significant associations between the percent signal change and cortical thickness in the previous analysis of the early blind group). The table presents estimates for contrasts in cortical regions which showed a significant interaction between group and cortical thickness. Confidence intervals for the trends as a function of group and contrast.*

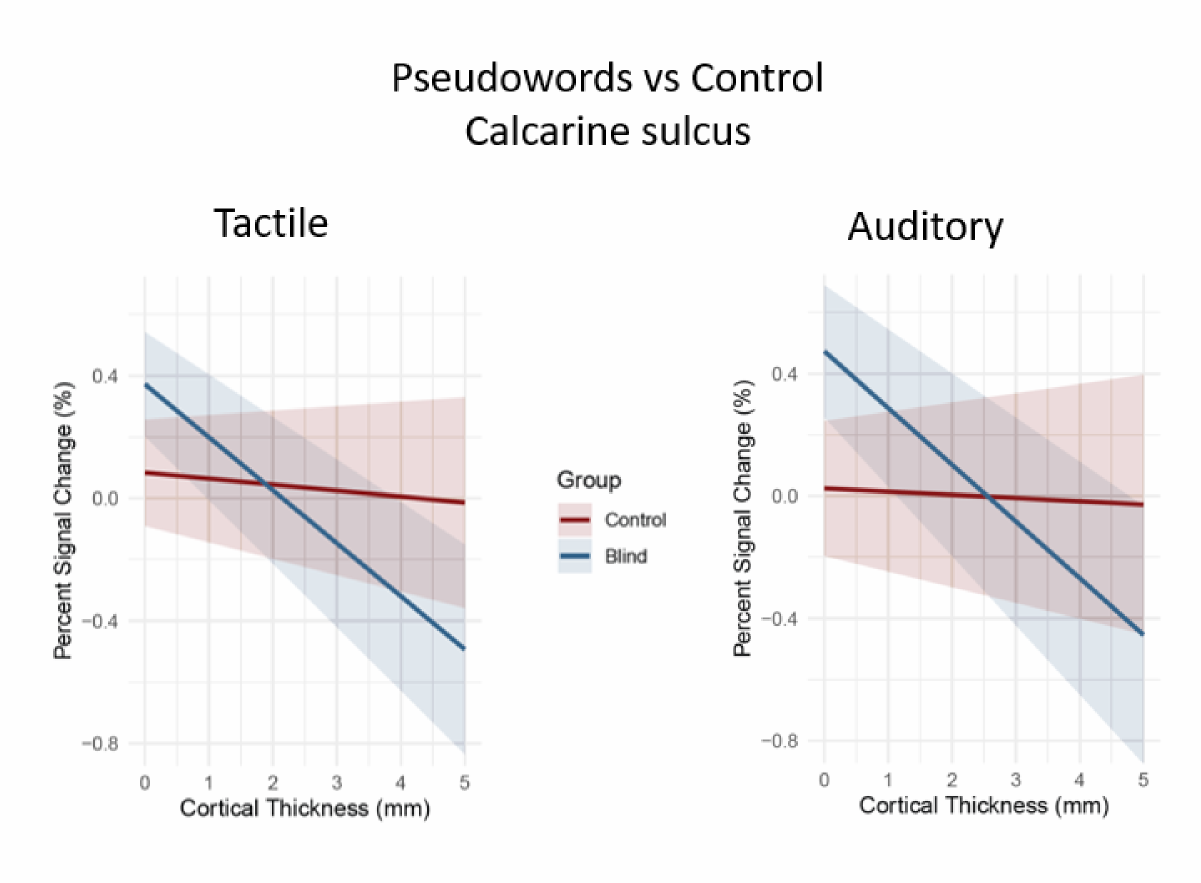


*Supplementary Figure 3. Cortical regions with significant interaction between cortical thickness and group (blind group n=25, control group n=25). The plots show population-averaged fitted curves of contrast- and ROI-specific linear mixed-effects models in which the main effect for cortical thickness was significant at the p<.05 level after a per-contrast Bonferroni correction. The ribbons represent confidence intervals.*
